# Amylin receptor subunit interactions are modulated by agonists and determine signaling

**DOI:** 10.1101/2024.10.09.617487

**Authors:** Sandra E. Gostynska, Jordan A. Karim, Bailee E. Ford, Peyton H. Gordon, Katie M. Babin, Asuka Inoue, Nevin A. Lambert, Augen A. Pioszak

## Abstract

Three amylin receptors (AMYRs) mediate the metabolic actions of the peptide hormone amylin and are drug targets for diabetes and obesity. AMY_1_R, AMY_2_R, and AMY_3_R are heterodimers consisting of the G protein-coupled calcitonin receptor (CTR) paired with a RAMP1, -2, or -3 accessory subunit, respectively, which increases amylin potency. Little is known about AMYR subunit interactions and their role in signaling. Here, we show that the AMYRs have distinct basal subunit equilibriums that are modulated by peptide agonists and determine the cAMP signaling phenotype. Using a novel biochemical assay that resolves the AMYR heterodimers and free subunits, we found that the AMY_1/2_R subunit equilibriums favored free CTR and RAMP1/2, and rat amylin and αCGRP agonists promoted subunit association. A stronger CTR-RAMP3 transmembrane domain interface yielded a more stable AMY_3_R, and human and salmon calcitonin agonists promoted AMY_3_R dissociation. Similar changes in subunit association-dissociation were observed in live cell membranes, and G protein coupling and cAMP signaling assays showed how these altered signaling. Our findings reveal regulation of heteromeric GPCR signaling through subunit interaction dynamics.

## Introduction

The peptide hormone amylin is a member of the calcitonin (CT)/calcitonin gene-related peptide (CGRP) family of peptides that is co-secreted with insulin from the pancreas in response to nutrient intake and acts at amylin receptors (AMYRs) in the brain to decrease food intake, slow gastric emptying, and decrease glucagon secretion (*1, 2*). The AMYRs are heterodimeric G protein-coupled receptors (GPCRs) that consist of a core class B GPCR subunit, the calcitonin receptor (CTR), in complex with a receptor activity-modifying protein (RAMP) accessory subunit that confers increased amylin potency (*3*). CTR co-expression with RAMP1, -2, or -3, gives rise to three amylin receptor subtypes, AMY_1_R, AMY_2_R, and AMY_3_R, respectively (*4–6*). These are drug targets for diabetes and obesity (*7*). The amylin analog pramlintide has been used as an insulin adjunct therapy for improved glycemic control (*8*). More recently, the long-acting, dual amylin and calcitonin receptor agonist (DACRA) cagrilintide was shown to promote substantial weight loss in clinical trials (*9–12*).

Little is known about the stability of AMYR heterodimers. If they are not stable complexes, then free subunits and heterodimers may co-exist at the cell surface, and the signaling phenotype may be a composite from both free CTR and AMYR heterodimers. While there are hints that this may occur (*13*), there has been no clear demonstration. Moreover, if AMYRs are transient heterodimers, it is possible that ligands could affect the subunit equilibrium. Recent cryo-EM structures of agonist-bound CTR- and AMYR-Gs complexes in detergent revealed differences in the CTR extracellular domain (ECD) orientation relative to the transmembrane domain (TMD) in the AMYRs engaged by human calcitonin (hCT) or the DACRA salmon calcitonin (sCT) as compared to rat amylin or αCGRP (*14, 15*). Interestingly, the sCT-AMY_1_R-Gs structure appeared to have a weakened CTR-RAMP1 interface at the ECD and top of the TMD. This hinted that some ligands may alter AMYR subunit interactions, but this remains unexplored. There are peptide drugs in development based on either the sCT or amylin backbones (*16–18*), highlighting the need to understand any differences in their actions at the AMYRs.

Here, we used a novel native PAGE mobility shift assay with detergent-solubilized AMYRs to resolve the heterodimers and free CTR and RAMP subunits. This revealed dramatic differences in the distribution of receptor species for the three AMYRs and changes upon ligand exposure, with the amylin/CGRP agonists promoting subunit association and the h/sCT agonists promoting dissociation in a subtype-selective manner. BRET assays demonstrated similar changes in subunit association-dissociation in live cell membranes, and signaling assays demonstrated coincident changes in G protein coupling and cAMP signaling. Together these studies revealed that the distinct subunit interaction dynamics of each AMYR and its modulation by peptide agonists determines the signaling phenotype.

## Results

### A native PAGE mobility shift assay with detergent solubilized AMYRs reveals the subunit distribution, its modulation by peptide agonists, and G protein coupling

We previously developed a biochemical native PAGE mobility shift assay for assessing agonist-dependent coupling of the engineered G protein surrogate miniGs (mGs) (*19*) to detergent-solubilized GPCRs (*20, 21*). We adapted the assay for the AMYRs and CTR to enable visualization of five species of interest: free RAMP, free CTR, CTR-RAMP heterodimer, ternary agonist-CTR- mGs complex, and quaternary agonist-CTR-RAMP-mGs complex (**Fig. 1A**). The CTR and RAMP constructs had a dual mCitrine-maltose binding protein (MBP) tag at their N-terminus and a native C-terminus. mCitrine permits visualization of each subunit by in-gel fluorescence and MBP improves expression and helps with electrophoretic separation of the various species. These constructs retained wt or near-wt pharmacology in cAMP signaling assays (**Fig. S1A-D**).

**Figure 1.**
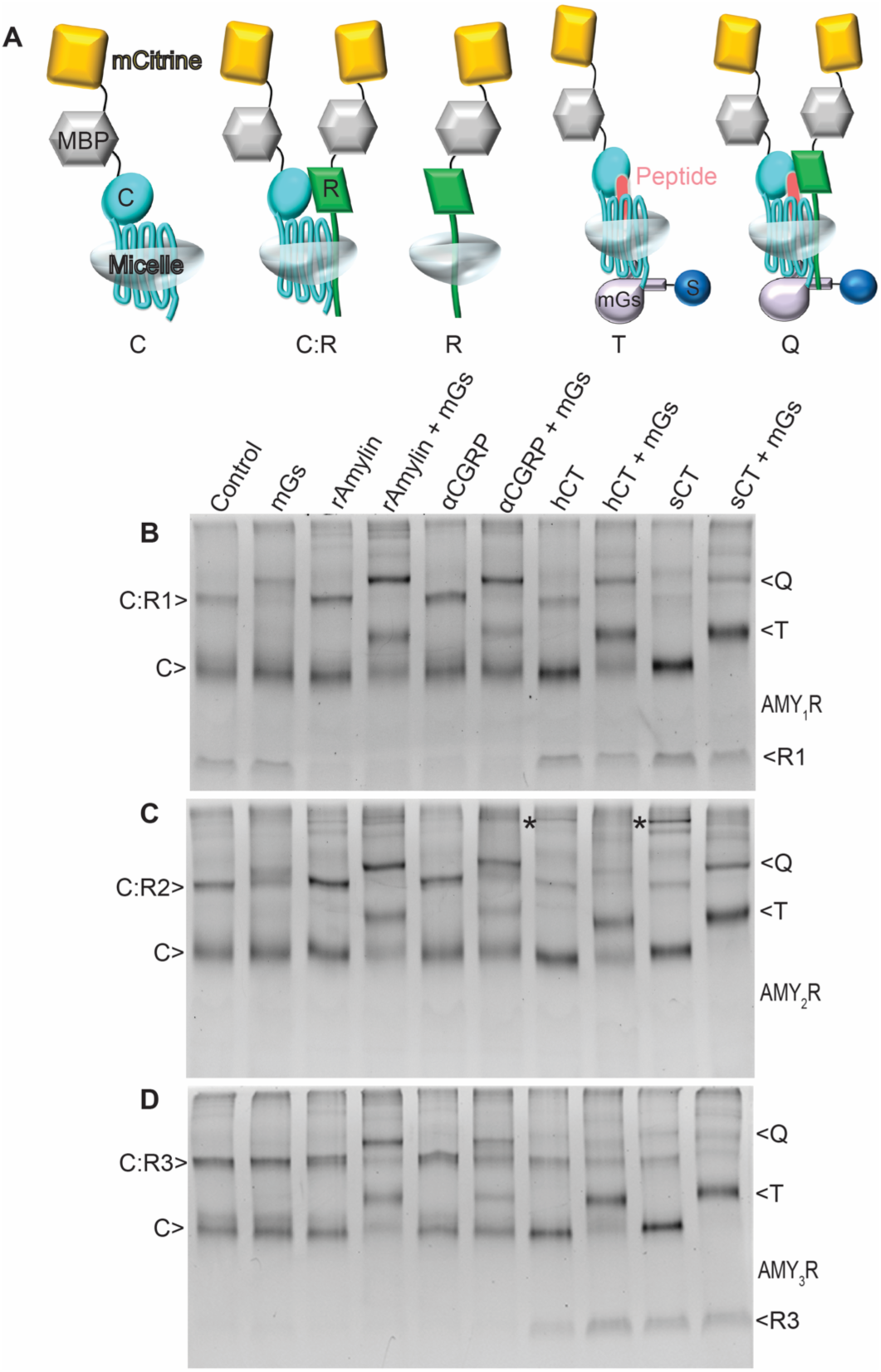
Visualization of free RAMP and CTR subunits and AMYR heterodimers in the absence or presence of agonists and miniGs by native PAGE mobility shift assay in detergent. **A**, Cartoon depicting the positions of the mCitrine-MBP tag on CTR (C) and RAMPs (R), and SUMO (S) tag on miniGs (mGs). **B-D,** Distribution of five molecular species C (CTR), C:R (CTR:RAMP), R (RAMP), T (ternary complex), Q (quaternary complex) formed in the presence or absence of 10 μM indicated peptide agonist and/or 50 μM purified SUMO-miniGs for co-expressed **B,** CTR and RAMP1 (AMY_1_R), **C,** CTR and RAMP2 (AMY_2_R), **D,** CTR and RAMP3 (AMY_3_R) in 7-10% gradient native gels. Representative gels for one of three replicates are shown with imaging for in-gel mCitrine fluorescence.

Using the tagged CTR and RAMP-expressing plasmids at CTR:RAMP transfection ratios of 1:1 or 1:2, free RAMP, free CTR, and the heterodimer were observed for each CTR-RAMP pairing (**Fig. S1E-G**). Higher order oligomeric species were also present, and these were particularly prominent with RAMP3. Decreasing the amount of the RAMP-expressing plasmid with CTR:RAMP ratios of 5:1, 10:1, or 20:1 significantly diminished the oligomers. We judged 5:1 and 10:1 to provide a reasonable balance of free CTR and heterodimer without significant oligomeric species. Under no conditions tested were we able to eliminate free CTR. Hereafter, the gel assay experiments used an 8:1 ratio of the CTR and RAMP constructs. First, we used qualitative single-point assays to examine the subunit distribution for each AMYR in the absence and presence of excess agonist (10 μM) and/or mGs (50 μM) (**Fig. 1B-D**). The agonists used were the 32-amino acid hCT and sCT and the 37-amino acid rat amylin (rAmy) and αCGRP. The CTR alone control is shown in Fig. S1h.

Co-expression of CTR and RAMP1 (AMY_1_R; **Fig. 1B**) yielded free RAMP1, free CTR, and the AMY_1_R heterodimer in the control lane (no agonist, no mGs). Addition of mGs alone shifted some of the heterodimer band to a higher molecular weight indicating some coupling to the AMY_1_R in the absence of agonist. Agonists alone did not yield shifts in band position, likely because the small peptides did not confer a sufficient change in mass. Nonetheless, agonist effects on the subunit distribution were evident because rAmy and αCGRP reduced free RAMP1 and increased the amount of the AMY_1_R heterodimer, consistent with their promoting subunit association. Together with mGs they yielded similar prominent quaternary complex bands that were more intense than with mGs alone, as well as ternary complex bands that were proportionally less favored. In contrast, hCT and sCT did not appreciably change the free RAMP1, and with mGs they yielded prominent ternary complexes and lesser amounts of the quaternary complexes.

Co-expression of CTR and RAMP2 (AMY_2_R; **Fig. 1C**) yielded free CTR and AMY_2_R heterodimer in the control lane and free RAMP2 was less apparent. Unfortunately, free RAMP2 runs as a broad smeary band (**Fig. S1F**), so this must be interpreted with caution. Nonetheless, rAmy appeared to increase the amount of AMY_2_R heterodimer, consistent with subunit association. rAmy and αCGRP supported both quaternary and ternary complex formation. hCT and sCT induced formation of a higher order oligomeric species of unknown molecular composition in the absence of mGs (* in Fig. 1c). These were absent in the presence of mGs where hCT yielded mostly ternary complex and sCT yielded both quaternary and ternary complexes.

Co-expression of CTR and RAMP3 (AMY_3_R; **Fig. 1D**) yielded free CTR and AMY_3_R heterodimer and little or no free RAMP3 in the control lane. The AMY_3_R heterodimer appeared to be unchanged with treatment of rAmy or αCGRP, but this might have reflected insufficient free RAMP3 for additional subunit association to occur. Both quaternary and ternary complexes were formed with rAmy and αCGPR in the presence of mGs. Notably, the two CT peptides increased free RAMP3, indicating their ability to dissociate the AMY_3_R heterodimer. Accordingly, they yielded mostly ternary complexes in the presence of mGs.

These results indicated that the native PAGE mobility shift assay is a robust biochemical assay for assessing AMYR subunit distribution, its modulation by ligands, and the coupling of each receptor species to G protein in detergent. Having established that AMYR heterodimers co-exist with free CTR and/or free RAMPs and that ligands can alter the subunit distribution, we next sought to quantitatively describe these effects.

### Native PAGE mobility shift assays with ligand titration quantitatively describe agonist modulation of subunit interactions

Agonist titration experiments were performed for each receptor in the presence of fixed, excess mGs (50 μM). The 4 agonists x 4 receptors gel matrix is shown in **Fig. S2A-P** and two key gels that show rAmy-promoted AMY_1_R association or hCT-promoted AMY_3_R dissociation are highlighted in **Fig. 2A,B**. Densitometry was used to quantitate the quaternary complex band for the three AMYRs and the ternary complex band for CTR alone. The former was normalized to the AMYR heterodimer and the latter to free CTR monomer bands in the control lanes. The potency (pEC_50_) and E_max_ values derived from the quaternary and ternary complex binding curves (**Fig. 2C-F**) are summarized in scatter plots (**Fig. 2G, H**) and **Table S1**. The potency can be thought of as measuring the apparent affinities of the agonists for the mGs-coupled receptors, although this is an over-simplification because we cannot saturate the receptor with mGs prior to agonist binding, and there are likely allosteric effects of mGs on agonist binding and vice versa. The E_max_ values provide insight into ligand modulation of subunit association/dissociation. For the quaternary complexes, E_max_ > 1 indicates subunit association and < 1 indicates dissociation. The changes in the free RAMP1/3 bands further illuminate the subunit interaction dynamics. These were quantitated and plotted in **Fig. 2I, J** with their pEC_50_ values shown in **Fig. 2K**.

**Figure 2.**
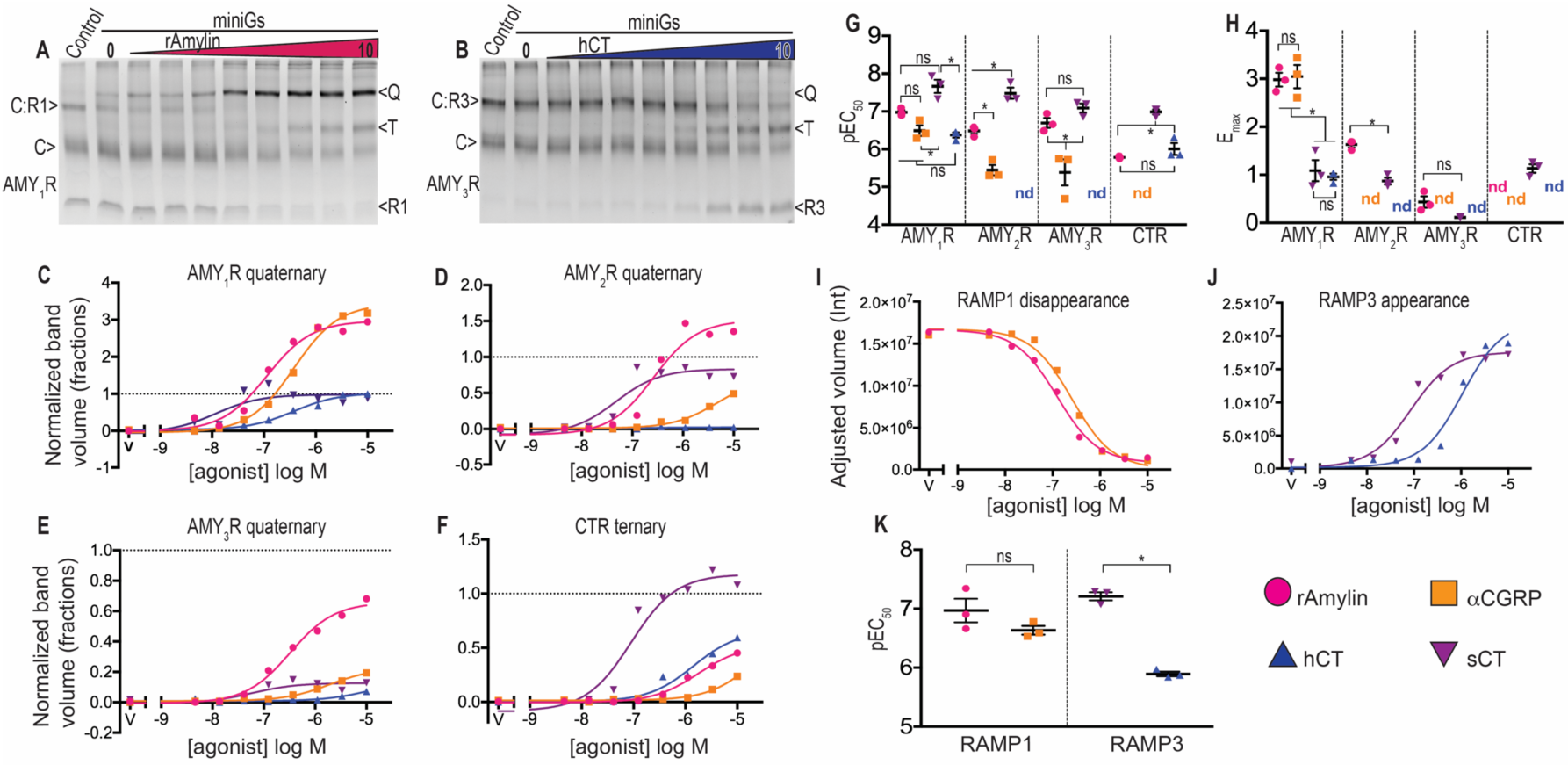
Densitometry quantitation of AMYR quaternary complexes, CTR ternary complexes, and free RAMP1/3 subunits from the agonist titration native PAGE experiments. **A, B**, Agonist titrations at detergent-solubilized CTR:RAMP heterodimers and free CTR in the presence of excess SUMO–miniGs by native PAGE mobility shift assay in 7-10% gradient native gels showing fluorescent bands for the co-expressed **A**, CTR and RAMP1 (AMY_1_R) and **B**, CTR and RAMP3 (AMY_3_R). **C-F**, Quantification of quaternary (Q) or ternary (T) complexes band appearance by densitometry at **A,** AMY_1_R, **B,** AMY_2_R, **C,** AMY_3_R and at **D,** CTR. For plots **C-F**, the quantitated gel band volumes were normalized to the heterodimer or free CTR bands in the control lane and represented in fractions. **G,** Scatter plot summarizing the replicate pEC_50_ ± SEM values for the agonist titration in c-f. **H**, Scatter plot summarizing the replicate E_max_ ± SEM values from C-F, nd = not determinable. See Supplementary Data Table S1 for a summary of the pEC_50_ and E_max_ values. **I,** Quantitation of free RAMP1 band disappearance or **J**, free RAMP3 band appearance. **K,** Scatter plot showing the replicate pEC_50_ ± SEM values for appearance of the free RAMP1; pEC_50_rAmy_ = 6.97±0.2 and pEC_50_αCGRP_ = 6.63±0.07 or disappearance of RAMP3 band; pEC_50_sCT_ = 7.21 ± 0.07 and pEC_50_hCT_ = 5.89 ± 0.04. The plots in **C-F**, and **I, J** show a representative from a single set of gels. Scatter plots in **G-H**, and **K** show the values from 3 independent replicates. Star indicates significance as compared with all other combinations determined by one-way ANOVA with Tukey’s post hoc test.

For AMY_1_R quaternary complex formation, rAmy and sCT were more potent than αCGRP and hCT (**Fig. 2C, G**), however, rAmy and αCGRP had E_max_ values 3-fold higher than the two CT peptides (**Fig. 2C, H**). The latter resulted from rAmy- and αCGRP-promoted CTR-RAMP1 association as evidenced by their E_max_ values of 3 and the decreased free RAMP1 with increasing agonist (**Fig. 2I**), which yielded pEC_50_ values in good agreement with those for quaternary complex formation (**Fig. 2K**). In contrast, the hCT and sCT E_max_ values of 1 suggested little or no effect on subunit association/dissociation.

For AMY_2_R, rAmy and sCT were more potent than αCGRP and hCT yielded little if any quaternary complex (**Fig. 2D, G**). The rAmy E_max_ was ∼1.5, consistent with it promoting association, whereas the sCT E_max_ was ∼1 and the αCGRP E_max_ could not be determined (**Fig. 2D, H**). hCT promoted loss of the AMY_2_R heterodimer, but it is unclear if this represented dissociation because of the lack of a visible free RAMP2 band (**Fig. S2G**).

For AMY_3_R, rAmy and sCT exhibited similar potencies that were stronger than αCGRP, and hCT yielded little if any quaternary complex (**Fig. 2E, G**). The rAmy E_max_ was below 1 (**Fig. 2E, H**), however, because not all the AMY_3_R heterodimer was shifted to quaternary complex at 10 μM agonist (**Fig. S2I**), this E_max_ was not rigorously defined, and it was approaching 1. The αCGRP and hCT E_max_ values could not be determined, whereas the sCT E_max_ was ∼0.1 (**Fig. 2E, H**). hCT and sCT promoted AMY_3_R dissociation as evidenced by the increase in free RAMP3 with increasing h/sCT (**Fig. 2J**). sCT was more potent than hCT for AMY_3_R dissociation and the sCT pEC_50_ was in good agreement with that obtained for quaternary complex formation (**Fig. 2K**).

For CTR alone, sCT was more potent than hCT and the other ligands (**Fig. 2F, G**). The sCT E_max_ was 1, whereas the E_max_ could not be determined for the other agonists due to their lower potencies (**Fig. 2H**). These results are consistent with prior studies of CTR pharmacology (*22, 23*).

We also performed miniGs titration experiments for each receptor in the presence of fixed, excess agonist (10 μM) (**Fig. S3A-U**). In this format, the potency can be thought of as measuring the apparent affinities of mGs for the agonist-occupied receptors, and this reflects agonist efficacy, at least in part, because mGs mimics the nucleotide-free state of Gs after GDP release (*19, 21*). While this format is not as enlightening for subunit association/dissociation, it still provided notable findings. First, it suggested that G protein binding has little effect on the AMY_1/3_R subunit interaction dynamics because the free RAMP1/3 bands did not change with mGs titration. Second, the higher order oligomers (*) induced by h/sCT in the absence of mGs with RAMP2 disappeared with increasing mGs (**Fig. S3G,H**), suggesting that G protein binding alters higher order AMY_2_R subunit interactions. Last, many of the mGs binding curves were shallow or biphasic (**Fig. S3Q- T; Table S2**). Although speculative, these may provide a molecular basis for the shallow/biphasic cAMP signaling concentration-response curves that have been reported for CTR and AMY_1/3_R (*23, 24*).

Overall, the ligand titration experiments indicated that AMYR subunit interactions are modulated by peptide agonists in a subtype-selective manner. rAmy and αCGRP promote AMY_1_R subunit association and h/sCT promote AMY_3_R dissociation. rAmy also promotes AMY_2_R formation, whereas h/sCT may induce higher order oligomers that are disfavored by G protein binding.

### A BRET assay for CTR-RAMP proximity on the cell surface supports agonist modulation of subunit interaction dynamics in cells

Next, we explored ligand modulation of CTR-RAMP association/dissociation for receptors on the surface of live cells. We developed a NanoBiT-based BRET assay (*25*) to monitor CTR-RAMP interactions in the plasma membrane of HEK293 cells (**Fig. 3A**). CTR was tagged at the N-terminus with the HiBiT peptide, and the three RAMPs were N-terminally tagged with mCitrine. Exogenous addition of the membrane-impermeable LgBiT reagent reconstitutes Nanoluciferase (*26*) only on cell-surface CTR. The tagged CTR and RAMP constructs retained wt pharmacology in a cAMP accumulation assay (**Fig. S4A, B**). Similar to the gel assays, we used a 10:1 CTR:RAMP transfection ratio for the BRET proximity assay. A robust BRET signal was observed in cells co-expressing HiBiT-CTR and mCitrine-RAMP1, and specificity was confirmed by the absence of BRET between HiBiT-β2-adrenergic receptor (AR) and mCitrine-RAMP1 (**Fig. S4C**).

**Figure 3.**
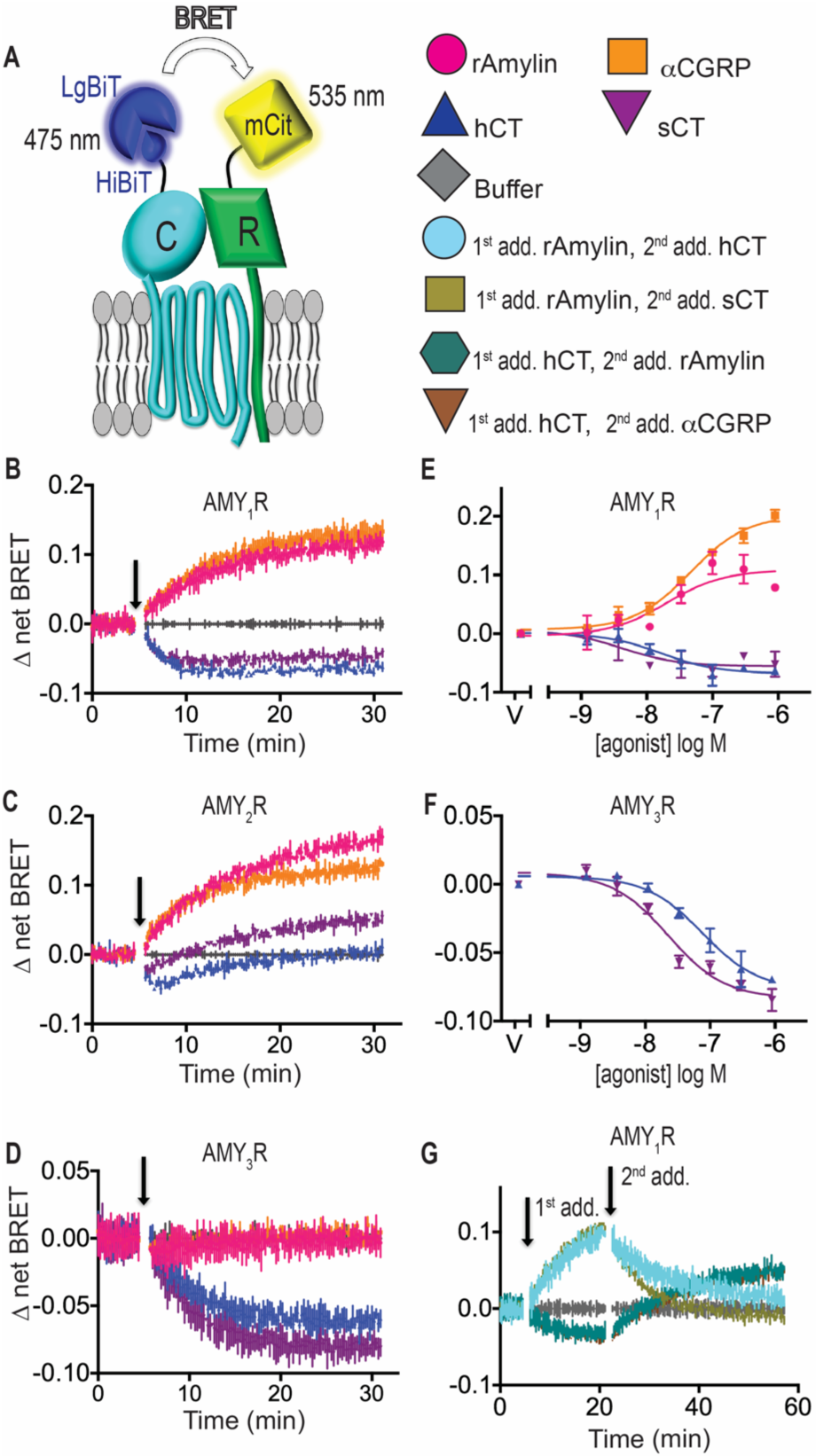
CTR:RAMP cell surface subunit proximity BRET assay in live HEK293 cells. **A**, Cartoon depicting the positions of the HiBiT tag at CTR and mCitrine tag at RAMP enabling signal measurement from CTR:RAMP heterodimers. **B-D**, Real-time kinetic CTR:RAMP proximity BRET was measured for co-expressed **B**, CTR and RAMP1 (AMY_1_R), **C**, CTR and RAMP2 (AMY_2_R), **D**, CTR and RAMP3 (AMY_3_R). Baseline was established for the first 5 min followed by agonist addition at 300 nM as indicated by the arrow. **E, F**, CTR:RAMP1 (AMY_1_R) and CTR:RAMP3 (AMY_3_R) in the endpoint concentration-response format. See Supplementary Data Table S3 for summary of pEC_50_ values. **G**, Real-time kinetic reversibility for co-expressed CTR:RAMP1 (AMY_1_R) with first addition of rAmy or hCT at 50 nM followed by second addition of 1 μM hCT/sCT or 1 μM rAmy/αCGRP respectively. All plots show a representative of three independent experiments at room temperature with duplicate technical replicates. Error bars show the SD for technical replicates.

Proximity of the CTR and RAMP1, -2, or -3 subunits was monitored in real-time upon addition of 300 nM of each agonist at room temperature. For AMY_1_R, rAmy and αCGRP similarly increased the BRET signal with relatively slow kinetics, whereas hCT and sCT similarly decreased it (**Fig. 3B**). For AMY_2_R, rAmy and αCGRP increased the BRET signal and rAmy ultimately yielded a higher increase, whereas hCT and sCT showed a brief initial decrease followed by a gradual increase (**Fig. 3C**). For AMY_3_R, rAmy and αCGRP had no effect, whereas hCT and sCT both decreased the BRET signal (**Fig. 3D**). Similar results were obtained at 37°C, except that the magnitude of change was generally smaller, other than for sCT at AMY_2_R, and the kinetics were faster (**Fig. S4D, E, F**).

Concentration-response assays showed dose-dependency of the agonist effects on AMY_1/3_R subunit proximity (**Fig. 3E, F**). rAmy and αCGRP had similar EC_50_ values of ∼30 nM. hCT and sCT had EC_50_ values of 30 and 7 nM, respectively, at AMY_1_R, and 76 nM and 22 nM, respectively, at AMY_3_R (**Table S3**). Real-time “challenge” experiments with sequential additions of rAmy and h/sCT or vice versa showed reversibility of the effects at AMY_1_R (**Fig. 3G**). These results were consistent with the assay reporting agonist modulation of AMY_1_R subunit proximity at the cell surface. It is unlikely that receptor internalization affected the signal because the AMY_1_R exhibits little if any internalization in HEK293 cells (*27*). Nonetheless, we cannot exclude the possibility that the assay also reported heterodimer conformation because rAmy and αCGRP yield different CTR-RAMP ECD orientations as compared to hCT and sCT (*14, 15*). We also cannot rule out the possibility that the assay reported higher order oligomerization.

Despite these caveats, overall the subunit proximity assay results were consistent with rAmy and αCGRP promoting association of CTR and RAMP1/2, and hCT and sCT promoting partial or full dissociation of AMY_1_R and AMY_3_R. The h/sCT behavior at AMY_2_R may reflect an initial separation of donor and acceptor that was swamped out by formation of the higher order oligomers that were observed in the gel assays (**Fig. 1C and Fig. S3G, H**). The lack of an effect of rAmy and αCGRP at AMY_3_R may have been due to insufficient free RAMP3 available for association to occur, as observed in the gel assays. This suggested that AMY_3_R may be a more stable heterodimer than AMY_1/2_R.

## Cell-based BRET assays for G protein coupling reveal pharmacology of AMYR heterodimers

To define the pharmacology of the AMYR heterodimers in live cells without confounding signal from free CTR, we developed a G protein coupling assay (*28*) to assess recruitment of mGs to the AMYRs by BRET. The RAMP subunits were tagged with the donor Rluc8 at the C-terminus to measure the recruitment of venus acceptor-tagged mGs to the AMYR heterodimers (**Fig. 4A**). A CTR-Rluc8 construct was used to characterize CTR alone. CTR-Rluc8 retained wt pharmacology in cAMP accumulation assays, and the three AMYRs with the RAMP-Rluc8 subunits exhibited wt signaling potencies and slightly decreased maximal responses (**Fig. S5A-D**). The BRET assay was performed in a HEK293 3GKO cell line in which the Gs, Gq, and G12/13 families of alpha subunits were knocked out (*29*). In addition, co-expression of pertussis toxin catalytic subunit inhibited endogenous Gi proteins. Importantly, control experiments with these constructs and the Gs-coupled β2-AR showed that bystander BRET from recruitment of venus-mGs to untagged receptors co-existing in the membrane with Rluc8-tagged receptors was insignificant (**Fig. S5E- L**).

**Figure 4.**
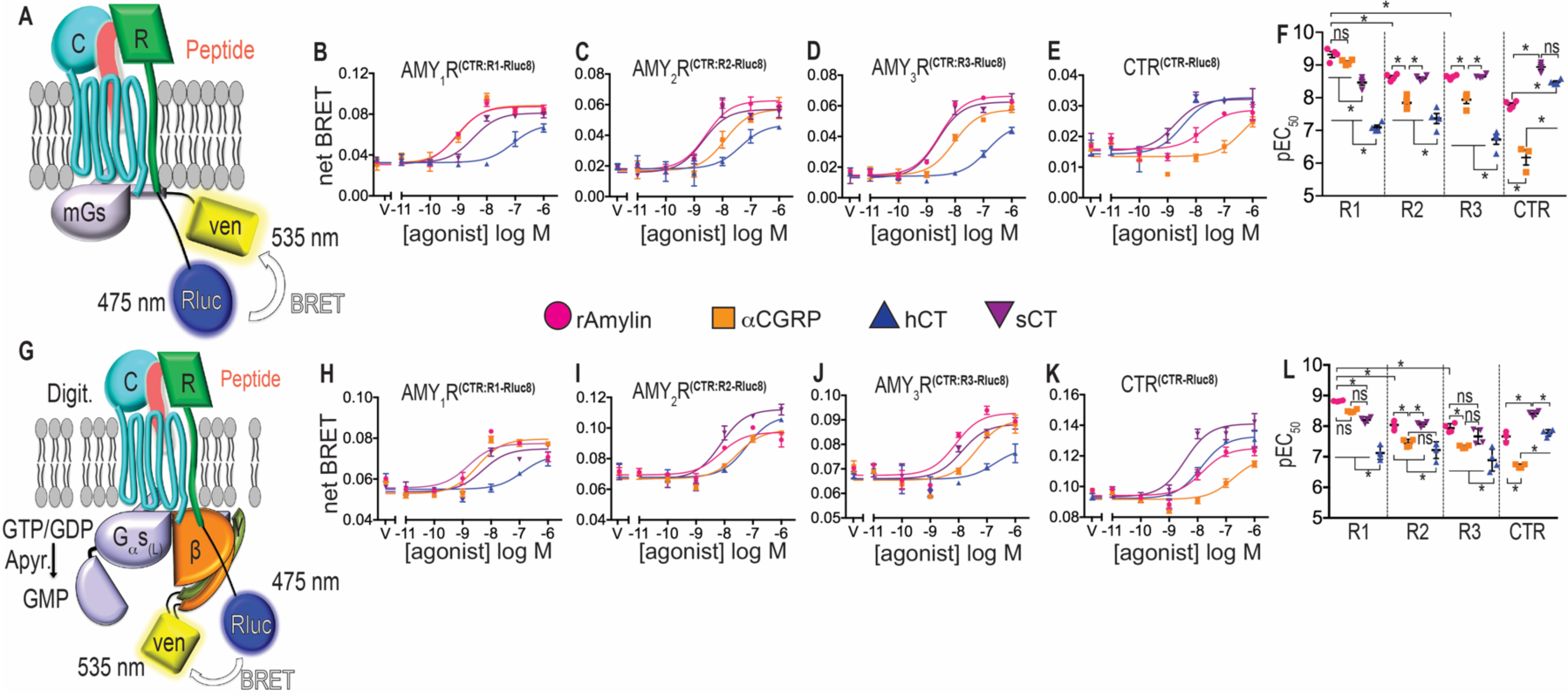
miniGs and heterotrimeric G protein coupling BRET assays for the AMY_1_R, AMY_2_R, AMY_3_R and CTR in live 3GKO HEK293 or permeabilized HEK293 cells. **A, G**, Cartoon depicting the positions of the Rluc8 donor and Venus acceptor used to isolate BRET signal from CTR:RAMP heterodimers in the two assay formats. **B-E**, Venus-miniGs recruitment to AMY_1_R (CTR:RAMP1-Rluc8), AMY_2_R (CTR:RAMP2-Rluc8), AMY_3_R (CTR:RAMP3-Rluc8) and CTR (CTR-Rluc8) in response to peptide agonist treatment; error bars show standard deviation for technical replicates. **F**, scatter plot summarizing pEC_50_ for each receptor– peptide combination as mean ± SEM from four independent replicates. **H-K**, Agonist mediated heterotrimeric G protein coupling to AMY_1_R (CTR:RAMP1-Rluc8), AMY_2_R (CTR:RAMP2-Rluc8), AMY_3_R (CTR:RAMP3-Rluc8) and CTR (CTR-Rluc8); error bars show standard deviation for technical replicates. **L,** Scatter plot summarizing pEC_50_ for each receptor–peptide combination as mean ± SEM from four independent replicates. Star indicates significance as compared with all other combinations determined by one-way ANOVA with Tukey’s post hoc test. See Supplementary Data Table 4 for summary of pEC_50_ values. Representative plots from a single replicate are shown for the concentration-response curves.

Concentration-response assays with the four agonists were used to characterize the mGs-coupling pharmacology of the three AMYRs and CTR (**Fig. 4B-E**). The agonist potencies are summarized in a scatter plot (**Fig. 4F**) and in **Table S4**. hCT was selective for CTR, at which it exhibited much stronger potency than at the three AMYRs. In contrast, sCT was a DACRA showing nearly equal strong potencies at all four receptor species, with perhaps a slight preference for CTR, although this did not reach statistical significance. rAmy and αCGRP both exhibited a selectivity profile of AMY_1_R > AMY_2_R = AMY_3_R > CTR. rAmy and αCGRP were equipotent at AMY_1_R, whereas rAmy was more potent than αCGRP at the other three receptor species, with αCGRP being particularly weak at CTR.

Next, we developed a version of the assay that used the Gs heterotrimer and wild-type alpha subunit. The enzyme apyrase was used to deplete GTP/GDP to trap the agonist-receptor-G protein complex in the nucleotide-free state (*30–32*). The Gγ subunits are tagged with the venus acceptor and the Rluc8 donor is located at the RAMP1/2/3 or CTR C-terminus to detect coupling to the three AMYRs or CTR, respectively (**Fig. 4G**). This assay, performed in wildtype HEK293 cells, gave similar results to the mGs-coupling assay (**Fig. 4H-L and Table S4**).

Recently, Keov *et al.* reported BRET G protein coupling assays specific for the AMYRs that also used mGs or Gs heterotrimer, but for the latter G protein turnover was allowed (*33*). The agonist potencies observed in their assays were weaker than in our assays, possibly due to their use of a shorter linker between the RAMP and the BRET donor that may have negatively affected receptor function. Their heterotrimer coupling assay yielded agonist selectivity profiles similar to our observations, however, their mGs coupling assay showed less selectivity of the agonists. The basis for the latter discrepancy in unclear, but it might reflect differences in methodology.

### CTR-RAMP interaction dynamics determines cAMP signaling phenotype

The results presented thus far suggested that co-expression of each RAMP with CTR may result in a distinct equilibrium distribution of molecular species on the cell surface. This could result from different AMYR stabilities. To test this, we used the native PAGE assay in a thermostability format (*21, 34*) to determine an empirical “melting temperature” (T_m_) for the three AMYRs in detergent (**Fig. 5A-C**). First, we calculated the fraction of the total CTR present as heterodimer for each AMYR incubated at 4°C. This revealed a higher proportion of CTR present as heterodimer with RAMP3 than RAMP1/2 (**Fig. 5D**). Quantitation of the disappearance of the heterodimer bands with increasing temperature revealed that the AMY_3_R (T_m_ 39°C) was significantly more stable than AMY_1_R and AMY_2_R (T_m_ 29°C) (**Fig. 5E, F**). This explains the differences in the subunit equilibriums in the basal state.

**Figure 5.**
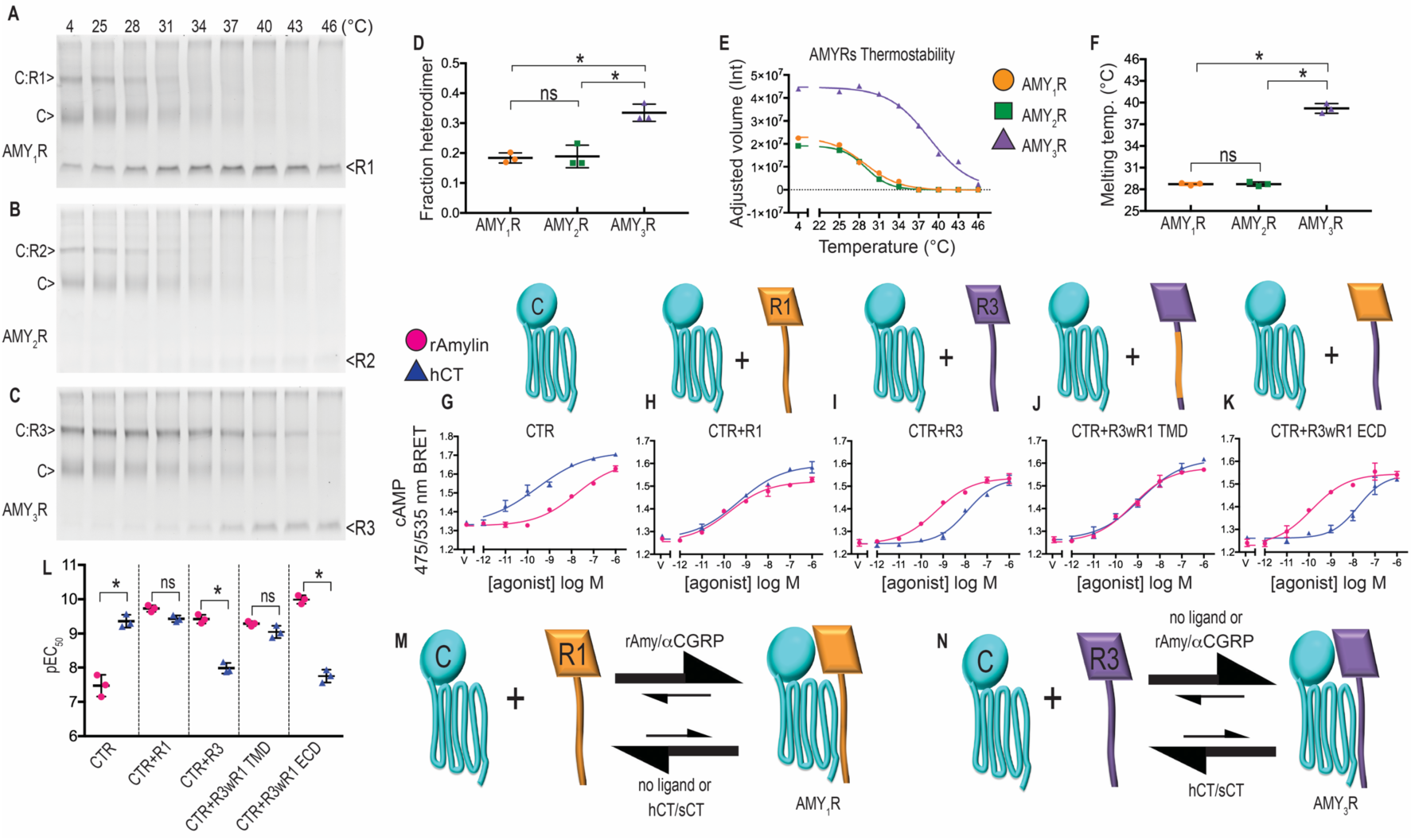
Thermostabilities of the AMYRs and cAMP signaling phenotypes in cells co-expressing CTR and wild-type or chimeric RAMPs. **A-C**, Stability of AMY_1_R (**A**), AMY_2_R (**B**), AMY_3_R (**C**) by native PAGE thermostability assay. Each gel is a representative unmodified image chosen out of three independent experiments and imaged for mCitrine fluorescence. **D**, Scatter plot summarizing fraction heterodimer for AMY_1_R, AMY_2_R, AMY_3_R from **A-C** at 4°C (lane 1) as mean ± SEM from three independent replicates. **E,** Quantitation of AMY_1_R, AMY_2_R, and AMY_3_R band disappearance by densitometry. Representative plot from a single replicate is shown. **E,** Scatter plot summarizing melting temperature (T_m_) for AMY_1_R T_m_ = 28.72°C ± 0.1, AMY_2_R T_m_ = 28.73°C ± 0.17, AMY_3_R T_m_ = 39.2°C ± 0.39 as mean ± SEM from three independent replicates. **G-K**, Concentration-response cAMP signaling phenotype in COS-7 cells transiently expressing **G**, CTR alone, **H**, CTR+RAMP1, **I**, CTR+RAMP3, **J**, CTR+RAMP3wR1 TMD, **K**, CTR+RAMP3wR1 ECD by cAMP CAMYEL biosensor BRET assay after 30 min agonist stimulation at 37°_C_. Error bars show standard deviation for technical replicates. Representative plots from a single replicate are shown. **L**, Scatter plot summarizing pEC_50_ for each receptor–peptide combination as mean ± SEM from three independent replicates. Star indicates significance as compared with all other combinations determined by one-way ANOVA with Tukey’s post hoc test. See Supplementary Data Table S5 for summary of pEC_50_ values. Model for **M**, CTR:RAMP1 and **N**, CTR:RAMP3 subunit equilibrium at the cell surface in the absence or presence of the indicated peptide agonists.

The different AMYR subunit equilibriums may explain the cAMP signaling phenotypes observed in cells co-expressing CTR and RAMPs (*4, 35*). We confirmed these in a BRET cAMP biosensor assay in COS-7 cells expressing CTR alone or CTR with RAMP1 or RAMP3 (**Fig. 5G-I, L and Table S5**). hCT was more potent than rAmy at CTR. Co-expression with RAMP1 enhanced rAmy potency and did not change hCT potency, whereas RAMP3 enhanced rAmy potency and decreased hCT potency. Because hCT is a much weaker agonist than rAmy at AMY_1_R (**Fig. 4B,H**), we reasoned that the strong cAMP signaling potency of hCT in cells co-expressing CTR and RAMP1 resulted from its activation of free CTR present in substantial amounts relative to AMY_1_R. In contrast, the swapped hCT and rAmy potencies observed with RAMP3 may result from less free CTR and more AMY_3_R on the cell surface due to the stronger RAMP3 interaction with CTR. In this case, the cAMP assay reports more AMY_3_R pharmacology, at which hCT is a weak agonist (**Fig. 4D,J**).

We hypothesized that the enhanced stability of the AMY_3_R resulted from the RAMP3 TMD interaction with the CTR TMD being stronger than that of RAMP1. If correct, co-expression of CTR with a chimeric RAMP3 having the RAMP1 TMD (“R3wR1 TMD”) should yield a decreased melting temperature and a cAMP signaling phenotype like that of CTR + RAMP1. These predictions were confirmed. In the thermostability assay, R3wR1 TMD yielded a decreased fraction heterodimer at 4°C (**Fig. S6A, C**) and a decreased melting temperature (**Fig. S6D, E**) that were similar to the RAMP1 heterodimer. In the cAMP assay, R3wR1 TMD yielded a phenotype like that of CTR + RAMP1 with rAmy and hCT being equipotent (**Fig. 5J, L**). Moreover, a chimeric RAMP3 in which the ECD was replaced with that of RAMP1 (“R3wR1 ECD”) retained a fraction heterodimer at 4°C (**Fig. S6B, C**) and thermostability (**Fig. S6D, E**) like that of RAMP3, and had a cAMP signaling phenotype similar to CTR + RAMP3 with rAmy being more potent than hCT (**Fig. 5K, L**).

## Discussion

Collectively, our data support a model in which the cAMP signaling phenotype in cells co-expressing CTR and RAMPs reflects the AMYR subunit equilibrium and its modulation by peptide agonists (**Fig. 5M, N**). In the basal state, the AMY_1_R equilibrium favors the free subunits due to a weaker CTR-RAMP1 transmembrane interface, whereas the AMY_3_R equilibrium favors the heterodimer due to the stronger CTR-RAMP3 transmembrane interface. The strong signaling potency of hCT in CTR + RAMP1 cells arises from its potent activation of free CTR, and its weak signaling potency in CTR + RAMP3 cells reflects its low potency at AMY_3_R and little free CTR. The strong signaling potency of rAmy in CTR + RAMP1 cells results from its promotion of subunit association to form AMY_1_R at which it is more potent than free CTR, plus its activation of the small fraction of pre-existing AMY_1_R. In CTR + RAMP3 cells, the strong rAmy signaling potency reflects its strong activation of pre-existing AMY_3_R. Our data strongly suggested that AMY_2_R is like AMY_1_R with its equilibrium favoring the free subunits and rAmy promoting their association.

rAmy and αCGRP were “associators” for AMY_1/2_R in the native PAGE and BRET subunit proximity assays. In contrast, hCT and sCT were “dissociators” for AMY_3_R in the native PAGE assay, but their effects on AMYR subunit interactions in cells appear to be more complex. The h/sCT-induced decrease in the BRET signal in the AMY_1_R subunit proximity assay may reflect partial dissociation of the top half of the heterodimer as hinted at in the sCT-AMY_1_R-Gs cryo-EM structure (*15*). This is consistent with the lack of h/sCT-induced AMY_1_R dissociation in the native PAGE assay and the strong potency of sCT at AMY_1_R in the BRET G protein coupling assays. For AMY_3_R, the h/sCT-induced decrease in the BRET signal in the subunit proximity assay may reflect partial and/or full dissociation. The strong potency of sCT at AMY_3_R in the BRET G protein coupling assays and the weak potency of hCT in the cAMP signaling assay for CTR + RAMP3 cells seem to suggest that they did not promote full dissociation. Interestingly, however, the hCT concentration response curve in CTR + RAMP3 cells was steeper than the other curves (**Fig. 5I**). This is consistent with some full AMY_3_R dissociation at higher hCT concentrations giving rise to free CTR at which it was more potent and efficacious. Regardless of the extent of their effects on subunit interactions in cells, the actions of h/sCT are clearly distinct from those of rAmy/αCGRP. h/sCT tend to favor dissociation, whereas rAmy/αCGRP promote association. The exception to this is the AMY_2_R, where h/sCT may induce higher order oligomerization.

Our results reveal a striking difference in the mechanisms by which the RAMPs modulate the pharmacology of CTR and the related CTR-like receptor (CLR). CLR-RAMP1 is the primary receptor for αCGRP, whereas CLR-RAMP2/3 are receptors for adrenomedullin (AM) peptides. These function in the neuronal, cardiovascular, and lymphatic systems (*36, 37*). Unlike CTR, CLR requires association with the RAMPs for transport to the cell surface (*38*) and all three CLR-RAMP heterodimers appear to be stable complexes (*21*). The RAMPs determine CLR ligand selectivity through distinct RAMP ECD contacts with CGRP and AM (*34, 39–41*) and allosteric effects on CLR (*3, 42*). While RAMP ECD contacts with amylin increase its affinity for the AMYRs (*15, 43, 44*), our results strongly suggest that a significant component of the mechanism of RAMP modulation of CTR pharmacology is their differential effects on the AMYR subunit equilibrium. These are mediated by the RAMP TMD and yield distinct AMYR subtypes, with AMY_3_R being more stable than AMY_1/2_R. The distinct cAMP signaling phenotypes reflect these differing basal subunit equilibriums and the way they are shifted by agonists.

Our findings have implications for AMYR biology and drug development. The distinct subunit interaction dynamics of the AMYRs suggest that AMY_3_R pre-exists on amylin target cells, whereas formation of appreciable amounts of AMY_1/2_R requires amylin-promoted subunit association. This might enable different kinetics of signaling responses between the AMY_3_R and AMY_1/2_R, or perhaps allow unique signaling events upon bringing RAMP1/2 together with CTR. The different, subtype-selective effects of sCT and rAmy on AMYR subunit equilibrium highlight the potential for diabetes/obesity drugs developed based on these two scaffolds to yield different signaling outputs and therapeutic outcomes. This will be important to consider in the interpretation of peptide efficacies in pre-clinical and clinical studies. Our results also have broader implications for RAMP complexes with numerous other GPCRs (*45–47*). We suggest that the dynamic CTR- RAMP complexes are a better model for these than the stable CLR-RAMP complexes. Subunit equilibrium considerations are likely to be relevant for RAMP modulation of other GPCRs, and the methods developed here may be of value for studying these complexes. The native PAGE method may also have broader utility for studying class A GPCRs, a large family with many members that can form dimers or higher order oligomers (*48, 49*).

## Methods

### Reagents

General chemicals and branched PEI were from Millipore-Sigma. Molecular biology enzymes were from New England Biolabs. LMNG and CHS were from Anatrace (Maumee, OH). Digitonin was from EMD Millipore. Linear PEI MAX (MW 40,000) was from Polysciences, Inc. LgBiT (N4018) and Nano-Glo® HiBiT Extracellular Substrate (furimazine) were from Promega, Madison, US (N3008). Coelenterazine h was from NanoLight. rAmylin, hαCGRP, hCT and sCT peptides were from Bachem (Torrance, CA). The lyophilized peptide powders were resuspended, quantitated, and stored as previously described (*34, 50*).

### Cell Culture

HEK293 (CRL 1573), HEK293S GnTI^-^ (CRL 3022) and COS-7 cells (CRL 1651) were from American Type Culture Collection (Manassas, VA). 3GKO HEK293 cell line with CRISPR/Cas9 mediated knockout of the *GNAS*, *GNAL*, *GNAQ*, *GNA11*, *GNA12* and *GNA13* genes was previously described (*29*). HEK293, 3GKO, and COS-7 cells were cultured in Dulbecco’s modified Eagle’s medium (with 4.5 g/l glucose and L-glutamine and 110 mg/L sodium pyruvate) from Gibco (11995–081) with 10% fetal bovine serum (Gibco 16000-044). HEK293S GnTI^-^ cells were grown in DMEM with 10% FBS and 1 X nonessential amino acids (NEAAs) from Lonza (Basel, Switzerland). Cells were grown at 37 °C, 5% CO_2_ in a humified incubator and passaged twice per week.

### Plasmids

The human CTR_(a)_, RAMP1-3, β2AR, Gαs(long), Gβ_1_, Gγ_2_ were used throughout and the pcDNA3.1-based expression plasmids for the wildtype proteins were from cdna.org. Plasmids encoding mVenus-miniGs, Venus-1–155-Gγ_2,_ Venus-156–239-Gβ_1_, and pRluc8-N1/β2AR-Rluc8 were previously described (*28, 30, 31*). The CAMYEL cAMP biosensor (*34, 50*) and pCAGGS/PTX-S plasmids were previously described.

Cloning of tagged CTR and RAMP constructs was performed by PCR/restriction digest/ligation methods. The native PAGE assay constructs used the pHLsec vector and its secretory peptide (*21*). CTR and RAMP1-3 were tagged at their N-terminus with monomeric citrine (mCitrine) and maltose binding protein (MBP) followed by 3C protease cleavage site. The constructs were as follows (with the CTR/RAMP amino acid residue numbers indicated): pHLsec/mCitrine-MBP- 3C-CTR.25–474 (pSG038), pHLsec/mCitrine-MBP-3C-RAMP1.24-148 (pJK040), pHLsec/mCitrine-MBP-3C-RAMP2.55-175 (pJK041), and pHLsec/mCitrine-MBP-3C- RAMP3.25–148 (pJK042). The mCitrine-MBP-3C tags were between AgeI and XmaI/EcoRV sites, CTR was from XmaI/EcoRV to XhoI, and the RAMPs were from XmaI/EcoRV to KpnI.

For G protein coupling BRET assays, CTR and RAMP1, -2 or -3 were C-terminally tagged with the luciferase Rluc8 and the natural receptor/RAMP secretory peptide was used. The constructs were as follows: pRluc8-N1/CTR.1-474-Rluc8 (pJK018), pRluc8-N1/RAMP1.1-148-linker_(27aa)_- Rluc8 (pPG027), pRluc8-N1/RAMP2.1-175-linker_(27aa)_-Rluc8 (pPG028), pRluc8-N1/RAMP3.1- 148-linker_(27aa)_-Rluc8 (pPG029). The CTR insert was cloned into the KpnI and AgeI restriction sites of pRluc8-N1 and the RAMPs were cloned into the EcoRI and AgeI sites.

The CTR:RAMP cell surface proximity BRET assay constructs used the pcDNA3.1(+) vector backbone and the signal peptide from the pHLsec vector. CTR and β2AR were N-terminally tagged with the HiBiT sequence (Promega) followed by an 8-amino acid linker, while RAMP1-3 were N-terminally tagged with mCitrine. The constructs were as follows: pcDNA3.1/HiBiT- CTR.25-474 (pJK030), pcDNA3.1/HiBiT-β2AR.2-413 (pJK031), pcDNA3.1/mCitrine- RAMP1.24-148 (pSG009), pcDNA3.1/mCitrine-RAMP2.55-175 (pSG010), and pcDNA3.1/mCitrine-RAMP3.25-148 (pSG011). For the CTR plasmid, the HiBiT tag was between AgeI and EcoRV sites followed by CTR ending with a XhoI site. The β2AR plasmid was similar except that HiBiT and β2AR were joined with EcoRV and ScaI sites that were destroyed upon ligation. For the three RAMP plasmids, mCitrine was between AgeI and XmaI/EcoRV sites followed by the RAMP ending with KpnI.

The RAMP3 with RAMP1 TMD or ECD chimeras were ordered as synthetic genes (Thermofisher geneart; not codon-optimized), which were cloned into the EcoRI and XhoI sites of pcDNA3.1(+) by Gibson Assembly for use in the cAMP signaling assay or in the pHLsec vector with mCitrine- MBP tags for the native PAGE assay. The plasmids encode untagged RAMP3.1-115-RAMP1.116- 142-RAMP3.143-148 (pKB046) or RAMP1.1-104-RAMP3.105-148 (pPG031) or their mCitrine-MBP tagged versions (pSG047 and pBF017).

The sequences of all plasmid inserts were confirmed by DNA sequencing by The University of Oklahoma Health Sciences Center Laboratory for Molecular Biology and Cytometry Research core facility. The plasmids were purified using a Macherey–Nagel midi kit according to the manufacturer’s instructions.

### Purified proteins

The H_6_-sumo-miniGs fusion protein and potato apyrase-H_6_ were expressed in bacteria and purified as previously described (*21, 51*).

### cAMP signaling assays

These were done in COS-7 cells and used the previously described CAMYEL BRET cAMP biosensor method (*50*) for validation of the native PAGE constructs and analysis of RAMP chimeras, or the LANCE cAMP accumulation method (*52*) for validation of the BRET assay constructs. In brief, cells were transiently transfected with 250 ng total DNA per well (96-well plates) using branched PEI. The plasmid ratios were CTR:RAMP:CAMYEL biosensor:pcDNA3.1(+) at 1:1:5:3 or CTR:CAMYEL biosensor:pcDNA3.1 at 1:5:4 for the biosensor assays or 1:1 CTR:RAMP or empty pcDNA3.1(+) for the LANCE assays. The cells were stimulated with agonist for 30 min at 37°C with addition of Coelenterazine h at 5 μM for the biosensor assay and the emissions at 475 and 535 nm were read in a PolarSTAR Omega plate reader. For the LANCE assay, the cells were stimulated for 15 min at 37°C in the presence of 1 mM IBMX followed by cell lysis and measurement of cAMP with the LANCE ultra cAMP detection kit (Perkin-Elmer or Revvity) according to the manufacturer’s instructions. The emissions at 620 nm and 665 nm were measured with the PolarSTAR plate reader. The concentration–response curves were fitted by nonlinear regression to a variable-slope dose– response stimulation equation in PRISM 7 (GraphPad Software, Inc., La Jolla,CA) to determine the pEC_50_.

### Native PAGE mobility shift assay for miniGs coupling

This assay was performed as previously described (*20, 21*), but with changes to the transfection procedure and preparation of the receptor-expressing cells. Here, we used whole adherent cells transfected in 6-well plates (other than for testing of transfection ratios, which used our original 48-well method). This eliminated well-to-well variation in transfection efficiency such that membrane preparations were not required to obtain high-quality data in the agonist or miniGs titration formats.

HEK293S GnTI^−^ cells were seeded at 1,200,000 cells per well into a 6-well plate in 2 mL/well growth media and grown overnight. The cells were transfected with 3 μg of DNA and 1.5:1 branched PEI:DNA ratio per well at 8:1:3 ratio of mCitrine-MBP-CTR:mCitrine-MBP-RAMP1- 3:pcDNA3.1 for the AMYRs or at 2:1 ratio of mCitrine-MBP-CTR:pcDNA3.1 for CTR alone. The DNA and PEI were combined in 100 μl per well of DMEM, incubated at room temperature for 10 min, and then valproic acid was added to 100 mM. The growth medium was removed by aspiration and replaced with 1.9 mL of pre-warmed (37°C) transfection media (1X DMEM with 2% FBS, 1X NEAA, 50 units/ml penicillin, 50 μg/ml streptomycin), and 0.1 mL per well of the transfection mix was added to cells. The cells were incubated for 72 hours at 30°C, 5% CO_2_. On the day of the assay, the medium was removed by aspiration, the cells were gently washed with 1X PBS, followed by aspiration. The cells were then harvested by trituration in 900 μL per well of ice-cold 1X harvest buffer (25 mM HEPES, pH 7.5, 140 mM NaCl, 10 mM KCl, 0.1 mM EGTA, 1X protease inhibitor (PI) tablet), and if necessary, cells from multiple wells were combined and transferred to a 5 mL tube on ice. The cells were gently pelleted by centrifugation for 5 min at 500 x *<u>g</u>* at 4°C, and the supernatant was discarded. The cells were resuspended in cold 1X reaction buffer (25 mM HEPES, pH 7.5, 140 mM NaCl, 10 mM KCl, 1 mM MgCl_2_, 2 mM CaCl_2_, 1X PI, 0.1 mg/mL FAF-BSA) to provide a cell suspension that was the source of the receptors for each assay format. For the qualitative single-point format, cells were resuspended in 70 μL/well and this was used as 7.62X, for the quantitative agonist titration format 53.2 μL/well gave 10X, and for the quantitative miniGs titration format 33.4 μL/well gave a 16X stock.

As in our prior work, the purified sumo-mGs fusion was pre-incubated in a GSH/GSSG redox buffer to ensure reduced cysteine residues before addition to the reactions (*20*). For the single-point qualitative assays, sumo-mGs, agonist, and cell suspension were combined as indicated in reaction buffer (25 mM HEPES, pH 7.5, 140 mM NaCl, 10 mM KCl, 1 mM MgCl_2_, 2 mM CaCl_2_, 1 X PI, 0.1 mg/mL FAF-BSA) and incubated 30 min on ice followed by LMNG/CHS solubilization for 2 hr at 4°C as described (*20*). The final reactions were 40 μL volume with 50 μM sumo-mGs, 10 μM agonist, and 1X cell suspension. For the quantitative agonist titration format, sumo-mGs and the cell suspension were combined in a master mix that was then added to an agonist dilution series followed by a 30 min incubation on ice and then 2 hr solubilization as above. Sumo-mGs was 50 μM in the final reactions and the agonists varied from 10 μM down in 3-fold increments. For the quantitative sumo-mGs titration format, agonist and the cell suspension were combined in a master mix that was then added to a sumo-mGs dilution series followed by a 30 min incubation on ice and then 2 hr solubilization as above. Agonist was 10 μM in the final reactions and the sumo-mGs varied from 75 μM down in 3-fold increments.

Native PAGE was performed as described (*20*), except that 7-10% polyacrylamide gradient resolving gels with a 5% stacking gel were used (made in-house). 20 μl of each reaction supernatant was loaded on the gels, which were run at 4°C for 3.5 hrs. Control samples with no agonist and no sumo-mGs were loaded in lane 1. The gels were imaged on a Chemidoc MP imager (BioRad) with 150 sec exposure using the ProQEmerald488 preset program. Densitometry for the titration experiments was performed as described (*20*) using the BioRad ImageLab software. The adjusted band volumes for the quaternary or ternary complexes were normalized to the heterodimer, or free CTR band volumes from the control lanes, respectively. The normalized adjusted volumes (fractions) were plotted against agonist or SUMO–mGs concentration in GraphPad Prism. Normalization was not used for quantitation of the free RAMP bands. The concentration–response curves were fitted by nonlinear regression to a fixed-slope dose–response stimulation equation in PRISM 7 (GraphPad Software, Inc., La Jolla,CA) for the agonist titrations to determine the pEC_50_ and E_max_. For the miniGs titrations, where biphasic binding was evident the data were fitted to a biphasic dose response equation in PRISM, constraining the bottom to 0 and slopes of the two phases to 1, or to a variable slope dose-response stimulation equation for the others. Fraction heterodimer was calculated as (heterodimer band volume/2)/((free CTR band volume + (heterodimer band volume/2)).

### Native PAGE thermostability assay

This assay was as described (*21, 34*), except that the cells were transfected by the above 6-well method with cells resuspended in 70 μL/well (7.62X) and the native PAGE conditions were as above. Densitometry was performed as above, and the adjusted heterodimer band volumes were plotted against temperature in GraphPad Prism with fitting of the “melt curves” to a variable slope logistic equation. The resulting IC_50_ or “melting temperature” (T_m_) is simply an empirical value that describes the breakdown of the AMYR heterodimers under the conditions of this assay.

### Live cell BRET assay for cell surface CTR-RAMP proximity

HEK293 cells were seeded at 20,000 cells/well and 100 μL/well in a 96-well white plate with clear bottom. After 24 hours, the cells were transiently transfected with 250 ng DNA per well and 1.5:1 branched PEI:DNA ratio. Cells were transfected with HiBiT-CTR, mCitrine-RAMP1, -2, or -3 and pcDNA3.1 at ratio of 10:1:14 and used for experiments 48 hrs later. Cells were washed with 1 X PBS and incubated for 30 min at room temperature or 37°C (as indicated) in 80 μL/well of 1X working buffer (25 mM Na-HEPES pH 7.4, 104 mM NaCl, 5 mM KCl, 1 mM KH_2_PO_4_, 1.2 mM MgSO_4_, 2 mM CaCl_2_, 1 mg/ml FAF-BSA, 5 mM glucose with 1/320 diluted LgBiT) in the dark. For the real-time kinetic format, 10 μL/well Furimazine was added to cells at 1/100 dilution (final) and incubated in the dark for 5 min. The emissions at 475 and 535 nm were read in a PolarSTAR Omega plate reader using a dual luminescence optic for 5 min to establish the baseline. 10 μL/well of the agonist was manually added to 300 nM final with reading for 30 min at room temperature or 37°C (as indicated). In the reversibility experiments, the first agonist addition was done manually to 50 nM with reading of emissions at 475 and 535 nm for 15 min followed by manual second addition of agonist to 1 μM and read for 30 min. Net BRET was calculated by subtracting 535/475 nm BRET ratio of cells expressing BRET donor with no acceptor from BRET ratio of cells expressing BRET donor and acceptor. Net BRET from a buffer only control was then subtracted to account for signal decay. Final delta BRET was plotted against time. For the endpoint concentration-response format, 10 μL/well Furimazine was added to cells at 1/100 dilution (final) and incubated in the dark for 10 min at room temperature. Peptide agonist serial dilutions were added as 10 μL/well and incubated for 20 min in the dark. Net BRET was plotted against agonist concentration and the concentration–response curves were fit by nonlinear regression to a fixed-slope dose–response stimulation equation in PRISM 7 (GraphPad Software, Inc., La Jolla,CA) to determine the pEC_50_.

### Live cell BRET assay for miniGs coupling

HEK293 3GKO cells were seeded at 500,000 cells/well and 2 mL/well in a 6-well plate. After 24 hrs the cells were transiently transfected using linear PEI MAX at 1.5:1 PEI:DNA ratio. Three μg total plasmid DNA was used per well with CTR, RAMP1- or -2-Rluc8, venus-miniGs, PTX-S1 and pcDNA3.1 at a ratio of 10:1:5:1:13 or at ratio of 20:1:10:2:27 for RAMP3-Rluc8. For CTR alone experiments, CTR-Rluc8, venus-miniGs, PTX-S1 and pcDNA3.1 were used in a 10:5:1:14 ratio. For miniGs bystander BRET control experiments with CTR, RAMP1-3-Rluc8, β2AR, venus-miniGs, PTX-S1, and pcDNA3.1 a ratio of 5:1:5:5:1:13 (RAMP1-, RAMP2-Rluc8) or a ratio of 10:1:10:10:2:27 (RAMP3-Rluc8) was used. The CTR-Rluc8 alone control experiments used CTR-Rluc8, β2AR, venus-miniGs PTX-S1 and pcDNA3.1 at a ratio of 5:5:5:1:14. After 48- h, the cells were washed once with 1 X PBS and were harvested by trituration with 900 μL/well of 1X working buffer (25 mM Na-HEPES pH 7.4, 104 mM NaCl, 5 mM KCl, 1 mM KH_2_PO_4_, 1.2 mM MgSO_4_, 2 mM CaCl_2_, 1 mg/ml fatty-acid-free bovine serum albumin (FAF-BSA), and 5 mM glucose). 50 μL of cell suspension was added per well in 96-well white plate with 25 μL/well of 4X agonist serial dilutions made in 1X working buffer. Coelenterazine h (25 μL/well) was added at 5 μM final concentration and incubated in the dark at room temperature. After 30 min of agonist exposure at room temperature, the emissions at 475 and 535 nm were read in a PolarSTAR Omega plate reader (BMG Labtech) using a dual luminescence optic. Net BRET was calculated by subtracting the BRET ratio of cells expressing BRET donor with no acceptor from the BRET ratio of cells expressing BRET donor and acceptor. The concentration–response curves were fitted by nonlinear regression to a fixed-slope dose–response stimulation equation in PRISM 7 (GraphPad Software, Inc., La Jolla,CA) to determine the pEC_50_.

### Permeabilized cell BRET assay for Gs heterotrimer coupling

This assay was performed as for the miniGs coupling assay, except for the following changes. Wt HEK293 cells were used, and they were transfected with CTR, RAMP1- 3-Rluc8, Gαs_long_, Venus- 1–155-Gγ_2_, Venus-155–239-Gβ_1_ at a ratio of 10:1:10:5:5 (RAMP1-, RAMP2-Rluc8) or 20:1:20:10:10 (RAMP3-Rluc8) for total of 3.1 μg DNA/well (RAMP1-, -RAMP2-Rluc8) or 3.05 μg DNA/well (RAMP3-Rluc8). Working buffer was replaced with 1X permeabilization buffer (140 mM KCl, 10 mM NaCl, 1 mM MgCl_2_, 0.1 mM EGTA, 20 mM Na-HEPES pH 7.2, 0.1 mg/mL FAF-BSA, 10 μg/mL digitonin). The agonist serial dilutions were in 1X permeabilization buffer supplemented with in-house purified apyrase (80 nM; 20 nM final).

### Statistical analysis

Experiments were done as ≥ three independent experiments on different days. For the cell-based BRET assays measuring miniGs or heterotrimeric G protein coupling and the cAMP biosensor assays, pEC_50_ values were compared using an ordinary one-way ANOVA with Tukey’s multiple comparisons test with statistical significance defined as *p* < 0.05. For agonist and SUMO–miniGs titration native PAGE assay experiments, comparison of pEC_50_ or E_max_ values for appearance of the quaternary/ternary complex band quantified by densitometry was done using an ordinary one-way ANOVA with Tukey’s multiple comparisons test. Statistical significance was defined as *p* <0.05. PRISM 7 was used for the statistical analyses.

## Supporting information

Gostynska_etal_2024supplemental

## Acknowledgements

This work was supported by grants from the National Institutes of Health (R01GM104251) and Oklahoma Center for the Advancement of Science and Technology (HR23-002) to AP, and NIH R35GM145284 to NL. AI was funded by the Society for the Promotion of Science (JP21H04791 and JP24K21281); the Japan Science and Technology Agency (JPMJFR215T and JPMJMS2023); and the Japan Agency for Medical Research and Development (JP22ama121038 and JP22zf0127007). We thank Dr. Dan Scott for kindly providing an mCitrine-encoding plasmid and Drs. Leo Tsiokas and Mohiuddin Ahmad for comments on the manuscript.

## Author Contributions

SG and AP conceived the study. SG, JK, PG, BF, and KB generated plasmid constructs. SG purified proteins. SG contributed to development and optimization of each of the assays and performed all assays. BF assisted SG on the native PAGE assays. JK and BF assisted with cAMP assays for tagged construct validation. JK contributed to the development and optimization of the native PAGE assay and KB contributed to optimization of the AMYR subunit BRET proximity assay. AI provided the HEK293 3GKO cell line and the PTX-expressing plasmid. NL conceived the BRET G protein heterotrimer coupling assay and performed initial feasibility experiments. SG, NL, and AP analyzed and interpreted the data. SG and AP contributed to visualization. AP acquired funding and supervised the project. SG and AP wrote the original draft, and the other authors were involved in reviewing and editing.

## Data availability

Data will be made available upon reasonable request to the corresponding author.

