## Supplementary material for "Amylin receptor subunit interactions are modulated by agonists and determine signaling": Gostynska_etal_2024supplemental

Sandra E. Gostynska *et al*.

* Augen A. Pioszak.

**This PDF file includes:**

Figs. S1 to S6
Tables S1 to S5

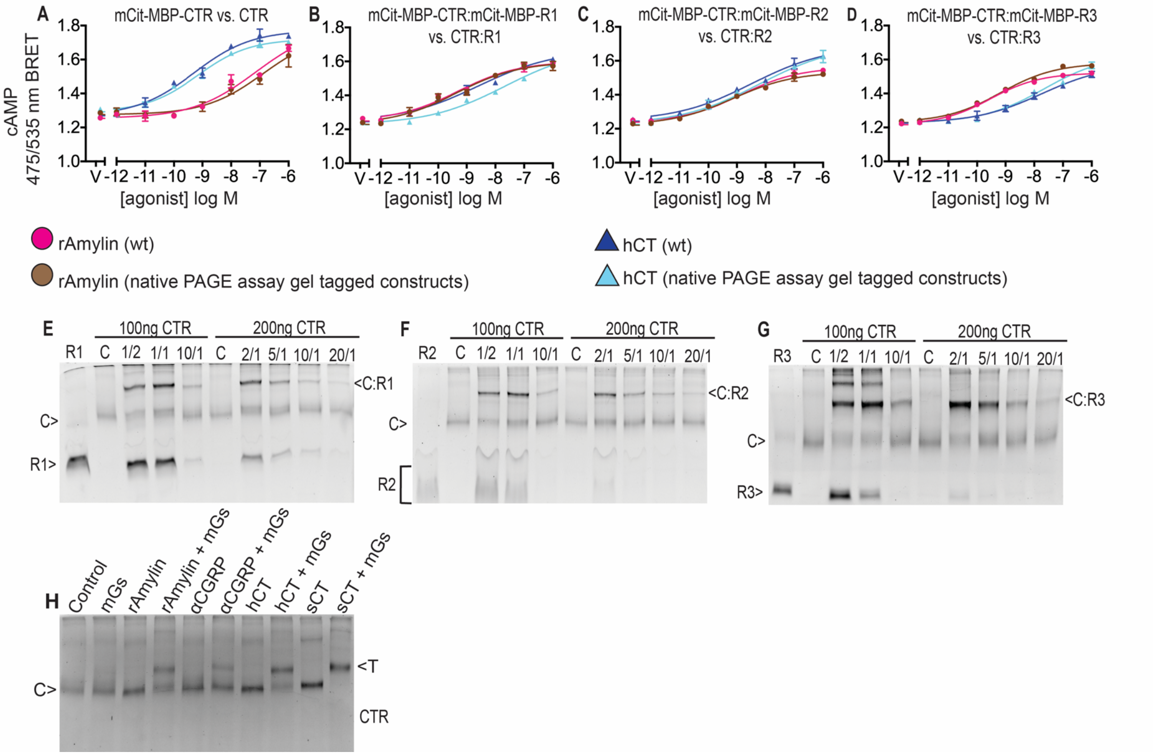


**Supplementary Figure S1. Validation of native PAGE assay constructs in cAMP CAMYEL biosensor assay and testing their transfection ratios in native PAGE assay. A-D,** Validation of native PAGE assay constructs in cAMP CAMYEL biosensor assay. **A,** mCitrine-MBP-CTR vs. wild type CTR, **B,** mCitrine-MBP-CTR:mCitrine-MBP-RAMP1 vs. wild type CTR:RAMP1, **C**, mCitrine-MBP-CTR:mCitrine-MBP-RAMP2 vs. wild type CTR:RAMP2, **D**, mCitrine-MBP-CTR:mCitrine-MBP-RAMP3 vs. wild type CTR:RAMP3. All plots show a representative of two independent experiments with duplicate technical replicates. Error bars have shown as mean ± SD for technical replicates. **E-G,** Testing transfection ratios of CTR and RAMP1-3 in native PAGE assay. HEK293S GnTI$^{-}$ cells were transiently co-transfected with CTR and RAMP at indicated ratios either at 100ng or 200 ng of CTR. **e**, mCitrine-MBP-CTR: mCitrine-MBP-RAMP1. **F**, mCitrine-MBP-CTR: mCitrine-MBP-RAMP2. **G,** mCitrine-MBP-CTR: mCitrine-MBP-RAMP3. 7%-9% hrCNE gradient gels were used. e-g, each gel is a representative unmodified image chosen out of 1-2 independent experiments. **H,** Distribution of species for CTR alone: C (CTR), T (ternary complex) formed in the absence or presence of 10 µM indicated peptide agonist and/or 50 µM purified SUMO-miniGs in 7-10% gradient native gels. Representative gel for one of three replicates is shown with imaging for in-gel mCitrine fluorescence.

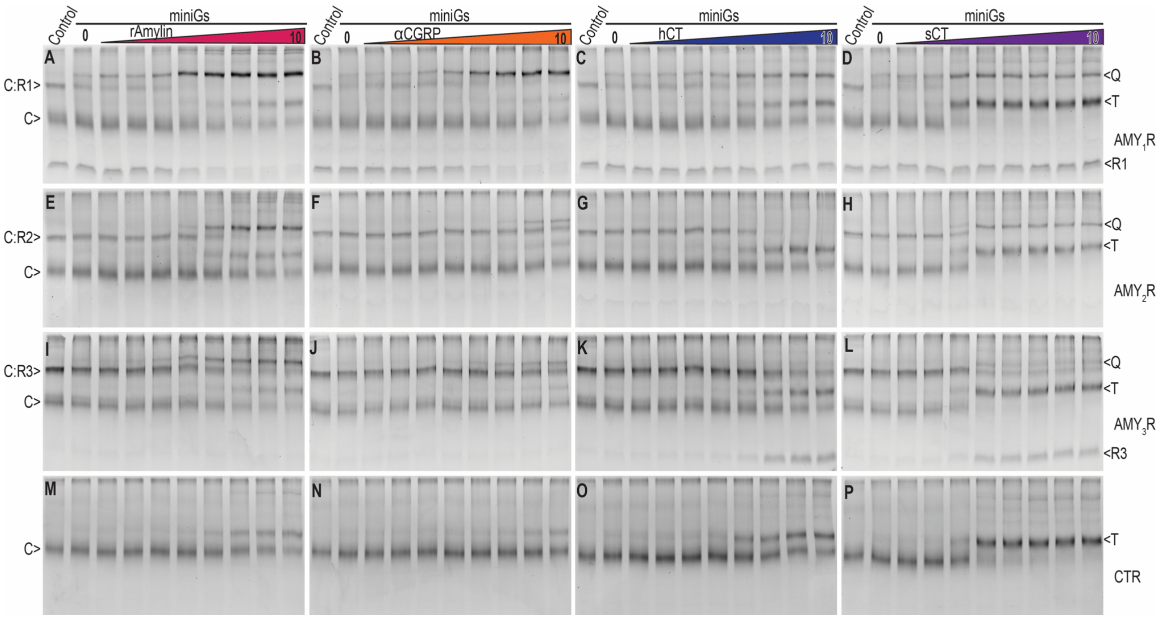


**Supplementary Figure S2. Agonist titrations at detergent-solubilized CTR·RAMP heterodimers and free CTR in the presence of excess SUMO–miniGs by native PAGE mobility shift assay. A-P**, 7-10% gradient native gels showing fluorescent bands corresponding to the co-expressed **A-D,** CTR and RAMP1 ($AMY_1R$), **E-H,** CTR and RAMP2 ($AMY_2R$), **I-L**, CTR and RAMP3 ($AMY_3R$), **M-P,** CTR alone. Quaternary (Q) or ternary (T) complexes formed by CTR:RAMP or CTR, respectively, in the presence of 50 μM SUMO–mGs and 3-fold serial dilutions of the indicated peptide agonists (10 μM highest). Control in lane 1 had no peptide or SUMO-miniGs treatment. Each gel is a representative unmodified image chosen out of three independent experiments with imaging for in-gel mCitrine fluorescence. Panels **A** and **K** are the same gels shown in **main figure 2 A, B** and are reproduced here for completeness.

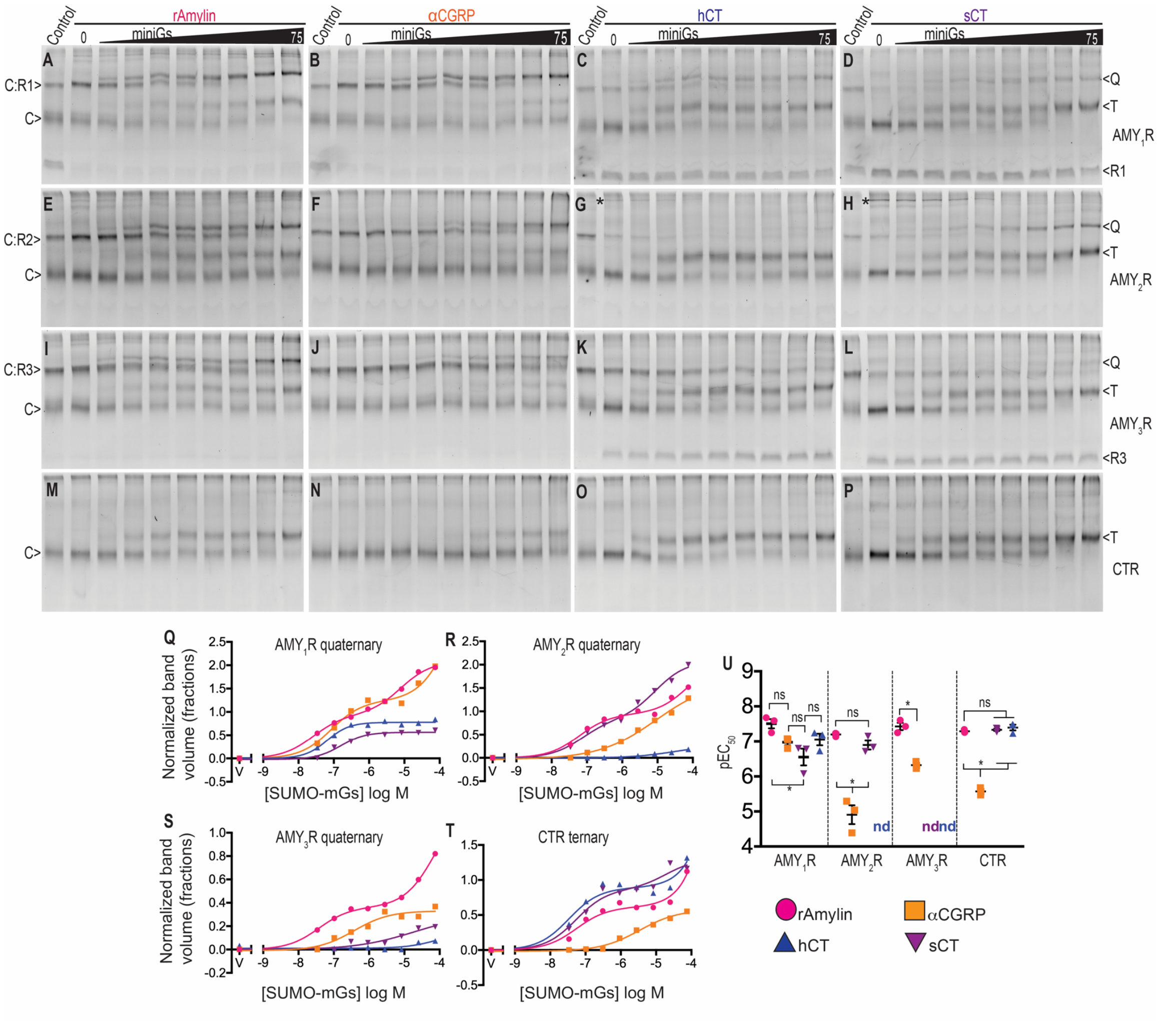


**Supplementary Figure S3. SUMO-miniGs titrations at detergent-solubilized CTR·RAMP heterodimers and free CTR in the presence of excess agonist by native PAGE mobility shift assay.** **A-P**, 7-10% gradient native gels showing fluorescent bands corresponding to the co-expressed **A-D,** CTR and RAMP1 ($AMY_1R$), **E-H,** CTR and RAMP2 ($AMY_2R$), **I-L**, CTR and RAMP3 ($AMY_3R$), **M-P,** CTR alone. Quaternary (Q) or ternary (T) complexes formed by CTR:RAMP or CTR, respectively, in the presence of the 10 μM indicated agonist and 3-fold serial dilutions of the SUMO-miniGs (75 μM highest). Control in lane 1 had no peptide or SUMO-miniGs treatment. Each gel is a representative unmodified image chosen out of three independent experiments with imaging for in-gel mCitrine fluorescence. **Q-T**, Quantitation of quaternary (Q) and ternary (T) complexes band appearance by densitometry at **Q,** $AMY_1R$ (from gels A-D), **R,** $AMY_2R$ (gels E-H), **S,** $AMY_3R$ (from gels I-L) and at **T,** CTR (from gels M-P). For panels **Q-T**, the quantitated gel band volumes were normalized to the heterodimer or free CTR bands in the control lane and represented in fractions. **U,** Scatter plot summarizing the replicate $pEC_{50} \pm$ SEM values for the SUMO-miniGs titration in q-t from three independent replicates, nd = not determinable. Star indicates significance as compared with all other combinations determined by one-way ANOVA with Tukey's post hoc test. See Supplementary Data Table S2 for a summary of the $pEC_{50}$ values.

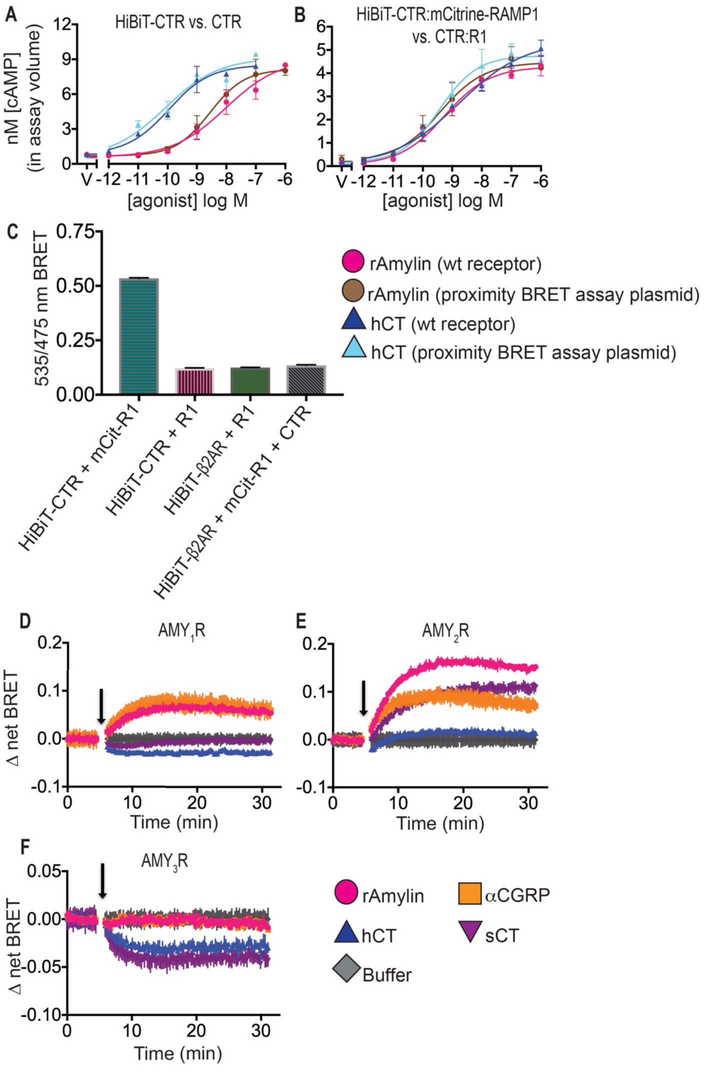


**Supplementary Figure S4. LANCE *ultra* cAMP end-point agonist concentration-response assay for validation of HiBiT-CTR and mCitrine-RAMP proximity BRET assay constructs and CTR:RAMP proximity BRET assay at 37°C. A**, HiBiT-CTR vs. wild type CTR, **B**, HiBiT-CTR:mCitrine-RAMP1 vs. wild type CTR:RAMP1 by LANCE *ultra* cAMP assay in COS-7 cells. **C**, End-point CTR:RAMP1 proximity assay in live HEK293 co-transfected with indicated constructs. Error bars are shown as mean ± SD for technical replicates. **D-F**, Real-time kinetic CTR:RAMP proximity BRET assay at 37°C in HEK293 cells for co-expressed **D,** CTR and RAMP1 ($AMY_1R$), **E,** CTR and RAMP2 ($AMY_2R$), **F,** CTR and RAMP3 ($AMY_3R$). Baseline was established for the first 5 min followed by agonist addition at 300 nM as indicated by the arrow. Plots show a representative of two or three independent experiments with duplicate technical replicates. Error bars show the mean ± SD for technical replicates.

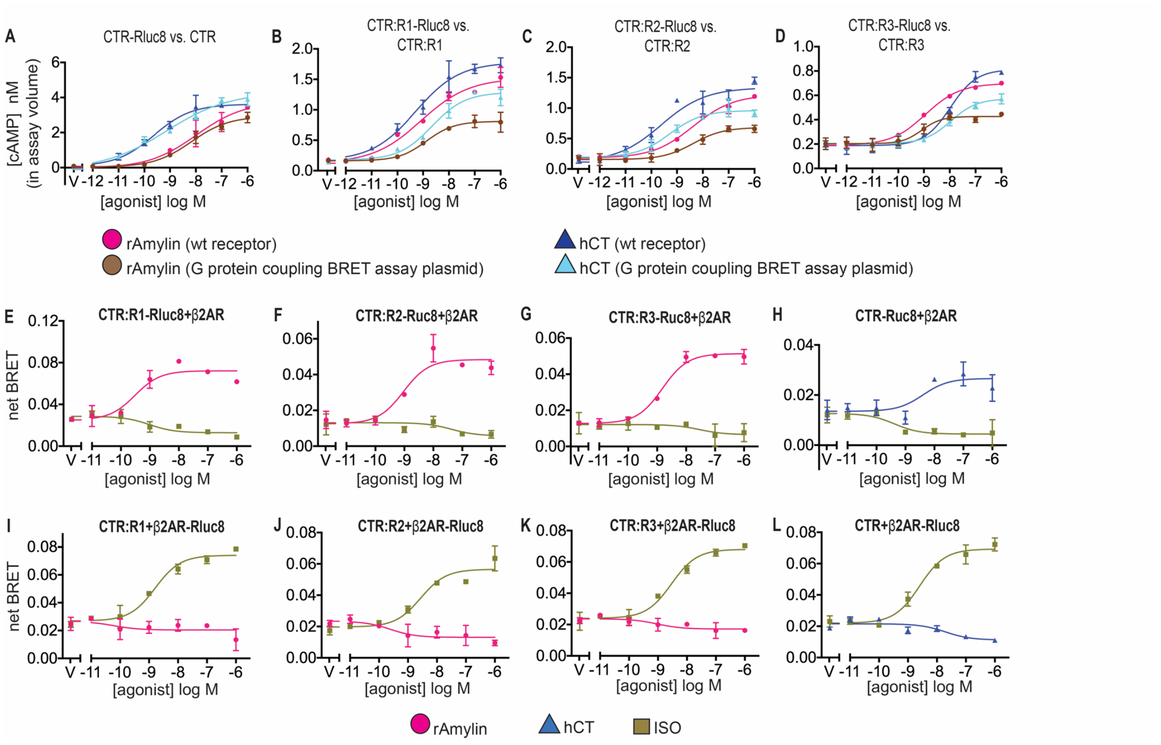


**Supplementary Figure S5. LANCE *ultra* cAMP end-point agonist concentration-response assay to validate CTR and RAMP-Rluc8 G protein coupling BRET assay constructs and venus-miniGs protein coupling bystander BRET assay controls. A**, CTR-Rluc8 vs. wild type CTR, **B**, CTR:RAMP1-Rluc8 vs. wild type CTR:RAMP1, **C**, CTR:RAMP2-Rluc8 vs. wild type CTR:RAMP2, **D**, CTR:RAMP3-Rluc8 vs. wild type CTR:RAMP3. All plots show a representative experiment with duplicate technical replicates. Error bars are shown as mean ± SD for technical replicates. **E-L**, venus-miniGs protein recruitment at **E**, CTR: RAMP1-Rluc8 (AMY$_1$R) with co-expressed β2AR, **F**, CTR:RAMP2-Rluc8 (AMY$_2$R) with co-expressed β2AR, **G**, CTR:RAMP3-Rluc8 (AMY$_3$R) with co-expressed β2AR, **H**, CTR-Rluc8 with co-expressed β2AR. **I-L**, venus-miniGs protein recruitment at **I,** CTR: RAMP1 (AMY$_1$R) with co-expressed β2AR-Rluc8, **J**, CTR:RAMP2 (AMY$_2$R) with co-expressed β2AR-Rluc8, **K,** CTR:RAMP3 (AMY$_3$R) with co-expressed β2AR-Rluc8, **L,** CTR with co-expressed β2AR-Rluc8. All plots show a representative of two independent experiments with duplicate technical replicates. Error bars are shown as mean ± SD for technical replicates.

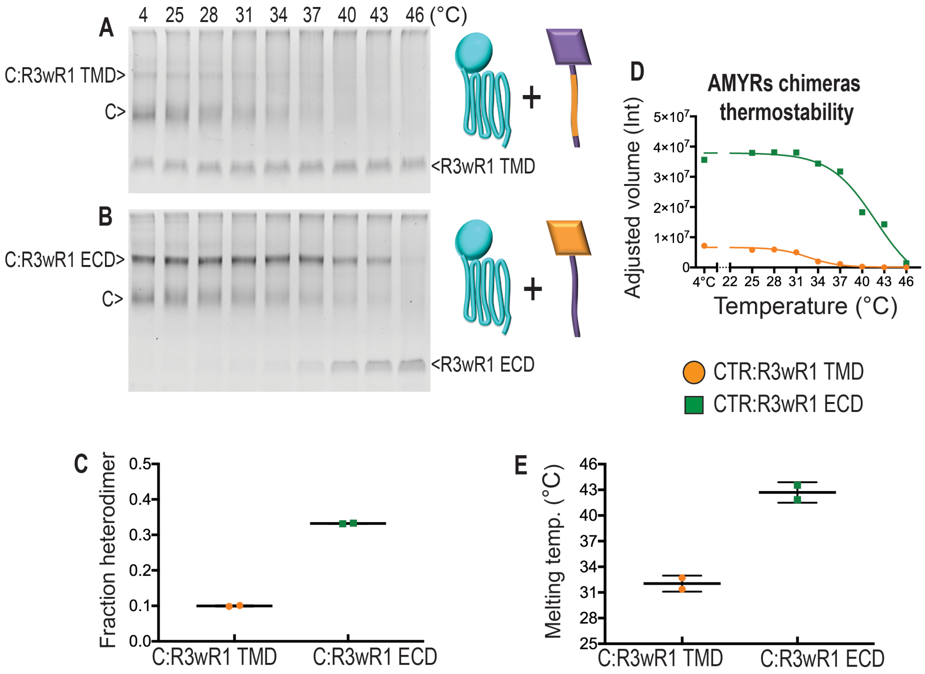


**Supplementary Figure S6. Thermostabilities of the AMYRs chimeras**. **A-B**, Stability of CTR:R3wR3_TMD chimera (**A**), CTR:R3wR1 ECD chimera (**B**) by native PAGE thermostability assay. Each gel is a representative unmodified image chosen out of two independent experiments and imaged for mCitrine fluorescence. **C**, Scatter plot summarizing fraction heterodimer for CTR:R3wR3_TMD chimera and CTR:R3wR3_ECD chimera from **A-B** at 4°C (lane 1) from two independent replicates. **D,** Quantitation of CTR:R3wR3_TMD chimera and CTR:R3wR3_ECD chimera band disappearance by densitometry. Representative plot from a single replicate is shown. **E,** Scatter plot summarizing melting temperature ($T_m$) for CTR:R3wR3_TMD $T_m$ = 32.04°C, CTR:R3wR3_ECD $T_m$ = 42.7°C. Error bars show mean with standard deviation for two independent replicates.

**Supplementary Data Table S1.** Summary of $pEC_{50}$ and $E_{max}$ values for agonist titration at CTR:R1-3 and CTR in the presence of excess SUMO- miniGs quantified by densitometry.

| Peptide agonist | **$pEC_{50}$ ± SEM** | | | | **$E_{max}$ ± SEM** | | | | N |
|---|---|---|---|---|---|---|---|---|---|
| | CTR:R1 ($AMY_1R$) | CTR:R2 ($AMY_2R$) | CTR:R3 ($AMY_3R$) | CTR | CTR:R1 ($AMY_1R$) | CTR:R2 ($AMY_2R$) | CTR:R3 ($AMY_3R$) | CTR | |
| rAmylin | 6.98 ± 0.06 | 6.49 ± 0.08 | 6.7 ± 0.13 | 5.78 ± 0.02 | 2.98 ± 0.14 | 1.63 ± 0.06 | 0.43 ± 0.12 | nd | 3 |
| αCGRP | 6.49 ± 0.14 | 5.44 ± 0.14 | 5.38 ± 0.35 | nd | 3.05 ± 0.24 | nd | nd | nd | 3 |
| sCT | 7.66 ± 0.18 | 7.49 ± 0.15 | 7.09 ± 0.12 | 6.98 ± 0.06 | 1.09 ± 0.22 | 0.87 ± 0.07 | 0.12 ± 0.01 | 1.13 ± 0.09 | 3 |
| hCT | 6.37 ± 0.07 | nd | nd | 6.01 ± 0.16 | 0.96 ± 0.07 | nd | nd | nd | 3 |

nd = not determinable

**Supplementary Data Table S2.** Summary of $pEC_{50}$ values ± standard error of mean for SUMO-miniGs titrations at CTR:RAMP1-3 and CTR in the presence of excess agonist quantified by densitometry.

| Peptide agonist | **$pEC_{50}$ ± SEM** | | | | N |
|---|---|---|---|---|---|
| | CTR:R1 ($AMY_1R$) | CTR:R2 ($AMY_2R$) | CTR:R3 ($AMY_3R$) | CTR | |
| rAmylin | 7.51 ± 0.13 | 7.2 ± 0.03 | 7.43 ± 0.10 | 7.29 ± 0.03 | 3 |
| αCGRP | 6.98 ± 0.09 | 4.91 ± 0.27 | 6.32 ± 0.06 | 5.57 ± 0.06 | 3 |
| sCT | 6.55 ± 0.24 | 6.9 ± 0.13 | nd | 7.33 ± 0.03 | 3 |
| hCT | 7.05 ± 0.16 | nd | nd | 7.4 ± 0.08 | 3 |

nd = not determinable
*For dose-response curves fitted to biphasic slope dose–response stimulation equation (Supplementary Data Figure. S3 Q-T) only first $pEC_{50}$ was reported.

**Supplementary Data Table S3.** Summary of $pEC_{50}$ values ± standard error of mean for CTR:RAMP1/3 proximity BRET assay dose-response endpoint in live HEK293 cells.

| Peptide agonist | **$pEC_{50}$ ± SEM** | | N |
|---|---|---|---|
| | $AMY_1R$ | $AMY_3R$ | |
| rAmylin | 7.84 ± 0.09 | - | 3 |
| αCGRP | 7.43 ± 0.05 | - | 3 |
| sCT | 8.16 ± 0.17 | 7.7 ± 0.06 | 3 |
| hCT | 7.53 ± 0.14 | 7.19 ± 0.45 | 3 |

**Supplementary Data Table S4.** Summary of $pEC_{50}$ values for venus-miniGs protein coupling BRET assay in live 3GKO cells and for heterotrimeric G protein coupling BRET assay in permeabilized HEK293 cell.

| Peptide agonist | **Venus-miniGs protein coupling BRET assay in live 3GKO cells $pEC_{50}$ ± SEM** | | | | **Heterotrimeric G protein coupling BRET assay in permeabilized HEK293 cells $pEC_{50}$ ± SEM** | | | | N |
|---|---|---|---|---|---|---|---|---|---|
| | CTR:R1 ($AMY_1R$) | CTR:R2 ($AMY_2R$) | CTR:R3 ($AMY_3R$) | CTR | CTR:R1 ($AMY_1R$) | CTR:R2 ($AMY_2R$) | CTR:R3 ($AMY_3R$) | CTR | |
| rAmylin | 9.32 ± 0.09 | 8.62 ± 0.0 | 8.64 ± 0.03 | 7.78 ± 0.06 | 8.82 ± 0.01 | 8.04 ± 0.08 | 7.94 ± 0.07 | 7.67 ± 0.08 | n = 4 |
| αCGRP | 9.08 ± 0.04 | 7.84 ± 0.10 | 7.94 ± 0.11 | 6.17 ± 0.23 | 8.49 ± 0.02 | 7.45 ± 0.06 | 7.32 ± 0.02 | 6.7 ± 0.03 | n = 4 |
| sCT | 8.47 ± 0.08 | 8.6 ± 0.04 | 8.65 ± 0.03 | 8.94 ± 0.07 | 8.21 ± 0.05 | 8.06 ± 0.04 | 7.67 ± 0.12 | 8.41 ± 0.04 | n = 4 |
| hCT | 7.02 ± 0.07 | 7.37 ± 0.15 | 6.73 ± 0.15 | 8.48 ± 0.04 | 7.13 ± 0.11 | 7.22 ± 0.14 | 6.89 ± 0.18 | 7.78 ± 0.05 | n = 4 |

**Supplementary Data Table S5.** Summary of $pEC_{50}$ values for cAMP signaling CAMYEL dose-response assay at CTR, CTR + RAMP1, or -3, CTR+RAMP3wRAMP1 TMD and CTR+RAMP3wRAMP1 ECD in COS-7 cells.

| Peptide agonist | **$pEC_{50}$ ± SEM** | | | | | N |
|---|---|---|---|---|---|---|
| | CTR | CTR + R1 | CTR + R3 | CTR+RAMP3wRAMP1 TMD | CTR+RAMP3wRAMP1 ECD | |
| rAmylin | 7.46 ± 0.32 | 9.73 ± 0.05 | 9.42 ± 0.07 | 9.29 ± 0.04 | 9.99 ± 0.07 | 3 |
| hCT | 9.43 ± 0.14 | 9.43 ± 0.06 | 7.99 ± 0.09 | 9.05 ± 0.10 | 7.75 ± 0.11 | 3 |
